# Variola virus is as old as the earliest historical lines of evidence suggest

**DOI:** 10.1101/2022.06.28.498054

**Authors:** Diego Forni, Cristian Molteni, Rachele Cagliani, Mario Clerici, Manuela Sironi

**Affiliations:** IRCCS E. MEDEA, Bioinformatics, Bosisio Parini, Italy; University of Milan, Milan, Italy; Don C. Gnocchi Foundation ONLUS, IRCCS, Milan, Italy

**Author notes:** These authors equally contributed to this work (Diego Forni, Cristian Molteni). Correspondence: Diego Forni.

**Keywords:** variola virus, smallpox, molecular dating, population structure

## Abstract

Archaeovirology efforts provided a rich portrait of the evolutionary history of variola virus (VARV), the causative agent of smallpox. These studies revealed frequent viral lineage extinction and a relatively recent origin of VARV as a human pathogen (∼ 1,700 years ago, ya). This contrasts with historical records suggesting the presence of smallpox as early as 3,500 ya. By performing an analysis of ancestry components in modern, historic, and ancient VARV genomes, we unveil the progressive drifting of VARV lineages from a common ancestral population and we show that a small proportion of Viking-Age ancestry persisted until the 18^th^ century. After the split of the P-I and P-II lineages, which occurred before the onset of smallpox vaccination, the former (but not the latter) experienced a severe bottleneck. We suggest this was due to the distinct epidemiology of the two lineages, which display remarkably different disease severity. As for the emergence of VARV, we corrected time estimates by accounting for the time-dependent rate phenomenon. Using this approach, we estimate that VARV emerged earlier than 3,800 ya, thus supporting its presence in ancient societies, including Egypt, as pockmarked mummies suggest.

**Importance:** Now eradicated, smallpox was one of the most devastating human diseases, causing the death of at least 300 million people in the 20^th^ century. Humans were the only known host of variola virus, but the time-frame of its emergence in our species has been a matter of debate. Specifically, molecular dating suggested a relatively recent origin, whereas historical sources indicated the presence of VARV in ancient societies. By applying population genetic methods, we analyzed the ancestry components in modern and historic VARV genomes, and we found a progressive drifting of VARV lineages from a common ancestral population. By accounting for the common observation that rates of viral evolution scale negatively with the time-frame of measurement, we estimated that VARV emerged earlier than 3800 years ago. Thus, out data settle a controversy and provide novel insight into the origin and evolution of one of the most historically relevant human pathogens.

## OBSERVATION

Variola virus (VARV), the causative agent of smallpox, represented a major cause of death during human history (1). Intensification of the vaccination campaign during the 20th century led to smallpox eradication in 1980 (1). At that time, two modern VARV (mVARV) lineages were circulating, P-I and P-II, the latter including alastrim (or variola minor) strains, characterized by much lower severity (2).

The time-frame of VARV emergence has been a matter of debate. Whereas some historical sources suggest that the disease was present in Egypt and Asia as early as 3,500 years ago (3), molecular dating analyses indicated a more recent origin for the virus. Specifically, the sequencing of ancient VARV genomes (aVARV) revealed that viral strains that were transmitted during the Viking Age (600 to 1050 CE) went extinct (4). This was also the fate of viral lineages that were sequenced from historical remains of the 17^th^ and 18^th^ centuries (historic VARV, hVARV) (5, 6). Here, we analyzed available modern, historic, and ancient VARV genomes to infer ancestry components and to re-assess the time-frame of VARV emergence.

## Results and Discussion

Like all orthopoxviruses, VARV has a long (∼185 Kbp) double-stranded DNA genome. We thus generated an alignment of 56 VARV genomes (4 aVARV, 4 hVARV, and 48 mVARV) and we analyzed 4013 parsimony-informative markers. These were used to evaluate the strength of linkage disequilibrium, which resulted moderate, thus warranting the application of models to analyze ancestry and admixture patterns implemented in STRUCTURE (7). This program can identify distinct subpopulations (or clusters, K) that compose the overall population and assigns each genome to one or more of those clusters. We applied the linkage model with correlated allele frequencies (7, 8) and we used an ancestry prior that allows source populations to contribute deferentially to the pooled sample of individuals (9). The ΔK method (10) estimated the best number of subpopulations to be equal to 3 (Figure S1). Analysis of ancestry components clearly separated aVARV, P-I, and P-II genomes, with only evidence of minor subpopulation sharing in one P-I sequence (strain India-1967). Conversely, hVARV genomes showed the contribution of all ancestral populations, although the aVARV component was the least abundant (Figure 1A). Two samples with controversial sampling dates (11) clustered with P-I and P-II lineages, in accordance with phylogenetic analyses (5). By estimating the F parameter, which represents a measure of genetic differentiation between populations, the linkage model also provides information on the level of drift of each subpopulation from a hypothetical common ancestral population. In line with sampling dates, the lowest drift was observed for the aVARV component. However, P-II had intermediate drift between aVARV and P-I components (Figure 1B).

**Figure 1.**
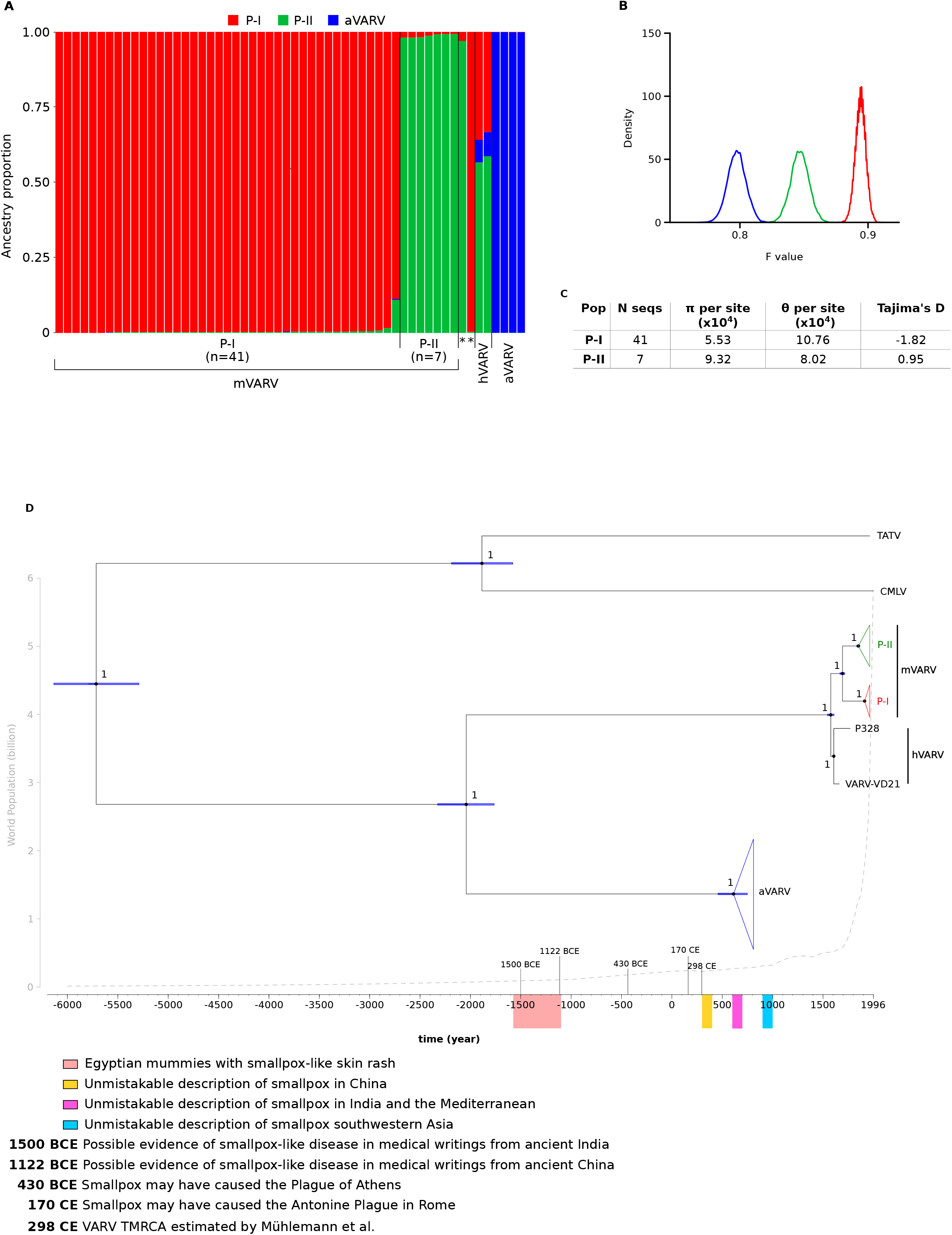
Population structure and molecular dating of VARV genomes. (A) Bar plot representing the proportion of ancestral population components. Each vertical line represents a VARV genome and it is colored by the proportion of sites that have been assigned to each population by STRUCTURE. Asterisks denote two samples (LT706529 and LT706528) with controversial dates (5, 11) (B) Distribution of F values for the three populations (colors as in (A)). (C) Nucleotide diversity estimators and Tajima’s D (14) for P-I and P-II clades. (D) Dated maximum credibility tree re-scaled after the TDRP correction. Branch lengths represent evolutionary time in years and a timescale grid is shown at the tree base, where some relevant historical events are highlighted (3). For each node, bars indicates 95% HPD intervals of node ages and the number indicates bootstrap support. World population size at different time points is reported as a gray dashed line (https://ourworldindata.org/).

Clearly, the history of VARV was characterized by frequent viral lineage extinctions, however these results portray a richer picture of VARV evolution. Identification of ancestry proportions unveils the progressive drifting of VARV lineages from a common ancestral population and indicates that a small fraction of Viking-age ancestry persisted until the 18^th^ century. Results herein also show and that the P-I lineage experienced a bottleneck that was more extreme than that of P-II (Figure 1B). This conclusion was also confirmed by the calculation of nucleotide diversity (12, 13) and Tajima’s D (14) (Figure 1C). However the small sample size for the P-II lineage suggests caution in the interpretation of these estimators.

One possible explanation for this finding is that, because of its generally lower virulence (2), the P-II lineage maintained an endemic, low-level transmission in some geographic areas, whereas the P-I lineage represented an epidemic, high-virulent lineage that was rapidly disposed of by control measures. Indeed, during the 20^th^ century, in areas where mild and severe smallpox forms were transmitted, the former persisted much longer in the population than the latter (1). Another, non-mutually exclusive possibility is that the two lineages were transmitted in regions where vaccination progressed with different pace and efficiency, thus resulting in different selective effects on circulating viruses.

We next re-assessed the time-frame of VARV emergence. Indeed, it is increasingly recognized that molecular dating can be affected by the time-dependent rate phenomenon (TDRP) - i.e., the negatively scaling of estimated rates of viral evolution with the time-frame of measurement (15, 16). We thus applied the recently developed PoW (prisoner of war) model (16) to convert units of branch lengths in the VARV phylogeny from substitutions/site to divergence time. Specifically, we used the median substitution rate estimated by Mühlemann and co-workers (4) (i.e. 5.16×10^−6^ s/s/y) and we applied the PoW conversion (16). To this purpose, we generated trees that also included taterapox virus (TATV, which infects gerbils) and camelpox virus (CMLV).

In line with the expectation under the PoW model, the TMRCA (time to most recent common ancestor) we estimated for the mVARV and hVARV clades (448 ya, 95% HPD: 416-482) and the mVARV TMRCA (332 ya, 95% HPD: 310-357) were very similar to previous estimates (Figure 1D, Table 1). However, the TMRCA of all VARV genomes was pushed back to ∼4,000 ya (95% HPD: 3790-4351), and we estimated that VARV split from the TATV/CMLV lineage ∼7,700 ya (Figure 1D). This latter finding does not imply that VARV entered human populations at that time, as the spill-over may have occurred at any time between the split from TATV/CMLV and the VARV TMRCA. Because TATV and CMLV evolved in non-human hosts with distinct generation times and biological features, the rate(s) of evolution in those species is unknown. We thus verified that the time estimates of the VARV phylogeny were not affected by the inclusion of the two animal viruses by repeating molecular dating for VARV sequences only. Time estimates were very similar to those obtained for the extended phylogeny (Table 1).

**Table 1.** Comparison between node ages estimated in this work and in previous ones.

| Estimate | This work<br>(with CMLV and<br>TATV) | This work<br>(VARV only) | Muhlemann et al. | Ferrari et al. | Duggan et al. |
| --- | --- | --- | --- | --- | --- |
| **TMRCa** | 5718 BCE | NA | NA | NA | NA |
| **TMRCa VARV** | 2045 BCE | 2767 BCE | 298 CE | NA | NA |
| **TMRCa**<br>**mVARV/hVARV** | 1574 CE | 1428 CE | 1534 CE | 1631/1677 CE | 1616 CE |
| **TMRCa PI/PII** | 1690 CE | 1600 CE | 1662 CE | 1789 CE | 1763 CE |
| **TMRCa PI** | 1910 CE | 1891 CE | ~1900 CE | 1892 CE | 1909 CE |
| **TMRCa PII** | 1847 CE | 1823 CE | ~1800 CE | 1866 CE | 1870 CE |

Previous estimates are from (4, 5, 6). NA: not available.

In the case of VARV, the time-frames we estimated are consistent with archaeological evidence of smallpox in ancient Egypt and with the presence of the disease in Asia in the second millennium BCE. A major criticism of the early existence of smallpox is the absence of any mention of it in written documents from ancient Egypt or in other early sources. However, the pattern of gene inactivation detected in aVARV genomes led to the suggestion (4, 17) that smallpox might have been a mild disease in the initial phases of human transmission. Thus, it might have gone unnoticed in early records. Of course, though, a number of other infectious diseases cause a rash similar to smallpox and only the sequencing of archeological specimens will provide information on which ancient societies were affected by the disease.

It should also be noticed that our study has limitations, which are inherent to the limited dataset of both modern and ancient VARV sequences, their biased geographic origin, and their passage history. Likewise, the long branches separating VARV, TATV, and CMLV underscore the extent of unsampled viral diversity for orthopoxviruses.

## Data availability

Materials and Methods are available as Supplementary Text S1.

## Competing interests

The authors declare no competing interests.

## Funding

This work was supported by the Italian Ministry of Health (“Ricerca Corrente 2022” to MS).

